# Proteomic and Transcriptomic Differences in Ischemic Stroke Patients with Atrial Fibrillation Versus Carotid Atherosclerosis

**DOI:** 10.64898/2026.07.02.736229

**Authors:** Daniela Renedo, Cyprien A. Rivier, Shufan Huo, Nanthiya Sujijantarat, Andrew Koo, Ryan Hebert, Richa Sharma, Kevin N Sheth, Dhasakumar S. Navaratnam, Lauren Sansing, Guido Falcone, Charles Matouk

## Abstract

**Background:** Ischemic stroke occurring in the setting of atrial fibrillation (AF) or carotid atherosclerosis (CA) may reflect distinct underlying biological processes. We integrated proteomic, transcriptomic, and genetic data from different sources to identify circulating proteins and molecular pathways associated with ischemic stroke in patients with AF versus CA.

**Methods:** We conducted a nested proteomic study within the UK Biobank comparing plasma protein levels among ischemic stroke patients with AF (n=539) and CA (n=127). Linear regression models were used to evaluate 2,923 proteins measured using the Olink Explore platform (false discovery rate [FDR] <0.05). In a separate Yale cohort, we evaluated expression of genes encoding identified proteins in thrombectomy clot single-cell RNA sequencing data from ischemic stroke patients with AF (n=7) or CA (n=7), including cell type–specific expression patterns. We then used summary statistics to perform 2-sample Mendelian randomization analyses using cis-protein quantitative trait loci to evaluate associations between genetically predicted levels of proteins identified in prior analyses and ischemic stroke subtypes. Exploratory pathway enrichment analyses were also performed.

**Results:** Twelve circulating proteins differed significantly between ischemic stroke patients with AF versus CA. AF was associated with higher levels of NTproBNP, NPPB, and ACP5, and lower levels of APCS, ANGPT2, PAMR1, PRCP, PROS1, LARP1, F7, F10, and LEO1 (all FDR<0.05). Clot transcriptomic analyses showed corresponding differential expression of ACP5, PRCP, LARP1, ANGPT2, and LEO1 across AF versus CA patients. Pathway analyses suggested enrichment of coagulation-related pathways among proteins associated with CA and natriuretic peptide signaling pathways among proteins associated with AF. Mendelian randomization analyses demonstrated associations between genetically predicted protein levels and ischemic stroke subtypes (AF or CA), including cardioembolic stroke for NTproBNPand ischemic stroke for ANGPT2, ACP5, APCS, and PAMR1.

**Conclusion:** **C**omplementary proteomic, transcriptomic, and genetic analyses identified differing molecular profiles among ischemic stroke patients with AF versus CA. These findings support established biomarkers, including NTproBNP and coagulation-related proteins, while identifying additional candidate pathways that may contribute to biological differences between these stroke-associated conditions. Further validation in clinically adjudicated and longitudinal cohorts is needed.

## Introduction

Embolic ischemic stroke is one of the leading causes of disability and death worldwide.(1) Among patients with ischemic stroke, atrial fibrillation (AF) and carotid atherosclerosis (CA) are common coexisting vascular conditions associated with differing recurrence risks, treatment strategies, and underlying biological processes.(2,3)However, distinguishing the relative contribution of these conditions to stroke pathophysiology can be challenging in clinical practice, particularly when conventional diagnostic evaluations yield inconclusive findings.(4)

Current biomarker strategies, including natriuretic peptides and coagulation-related proteins, have shown potential for improving understanding of stroke-associated biological pathways and identifying patients at increased risk of specific ischemic stroke phenotypes.(5) Nevertheless, most prior studies have relied on single-modality datasets, limiting the ability to evaluate whether observed associations are supported across complementary biological contexts.

The increasing availability of large-scale proteomic, transcriptomic, and genetic datasets provides an opportunity to examine convergent molecular signals associated with ischemic stroke in patients with AF or CA.(6) Although these data sources capture distinct biological states and are not integrated at the individual patient level, complementary analyses across orthogonal datasets may help identify candidate proteins and pathways consistently associated with stroke-related vascular phenotypes.

In this study, we used complementary proteomic, transcriptomic, and genetic approaches to evaluate molecular differences among ischemic stroke patients with AF versus CA. Specifically, we analyzed circulating plasma proteins in the UK Biobank, explored expression patterns of corresponding genes in thrombectomy clot single-cell RNA sequencing data, and performed Mendelian randomization analyses using genetically predicted protein levels and ischemic stroke phenotypes. Our objective was to identify candidate molecular pathways differentially enriched in ischemic stroke occurring in the setting of AF or CA and to generate hypotheses for future mechanistic and translational studies.

## Methods

### Study Design

We conducted complementary proteomic, transcriptomic, and genetic analyses across independent datasets to evaluate molecular differences among ischemic stroke patients with AF or CA (**Figure 1**).

**Figure 1.**
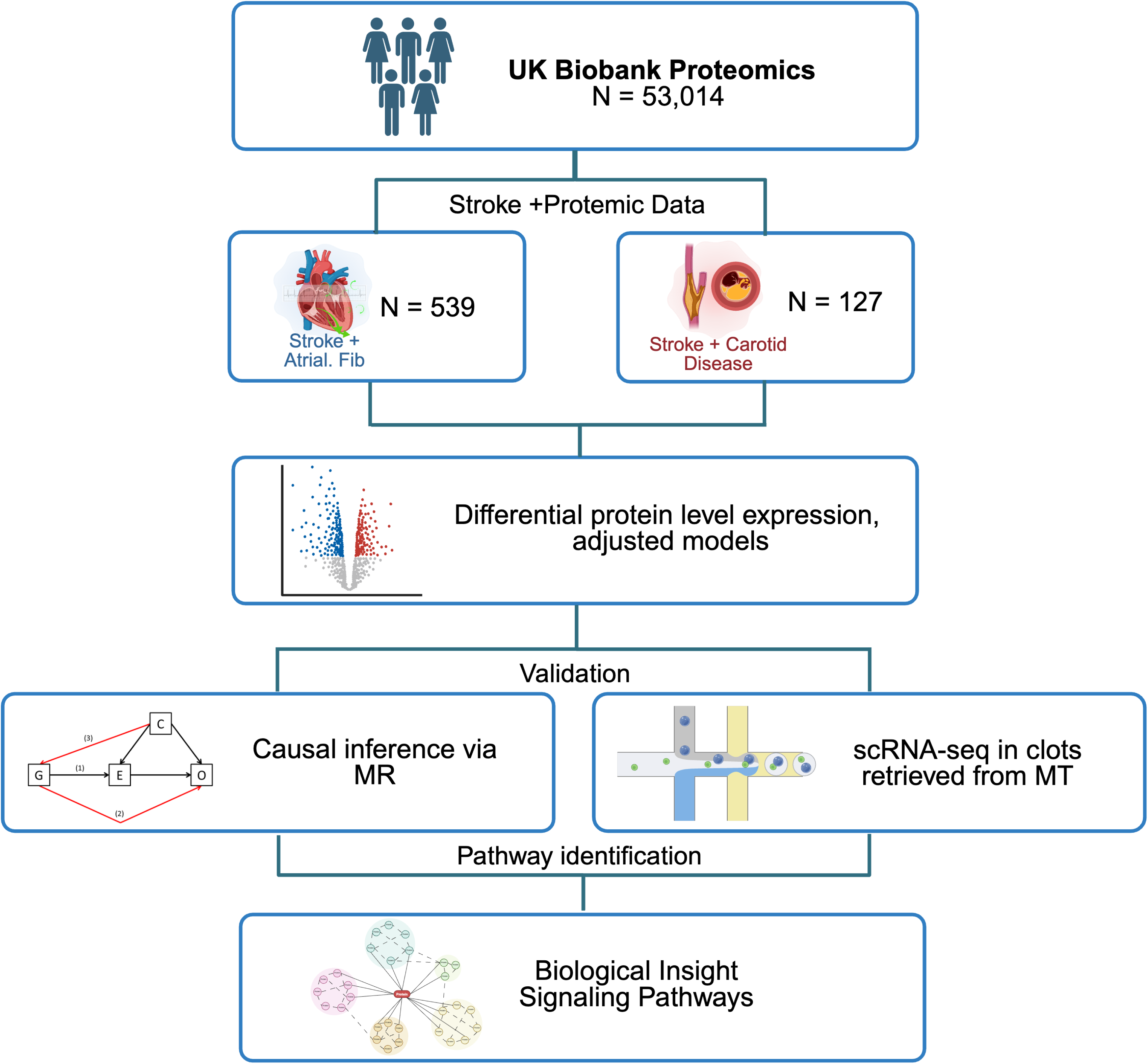
Study design and analytical workflow. UK Biobank participants with proteomic data (N = 53,014) were screened to identify patients with ischemic stroke and either atrial fibrillation (n = 539) or carotid artery disease (n = 127). Differential protein expression analyses using adjusted regression models were followed by Mendelian randomization and exploration analysis in single-cell RNA sequencing of thrombectomy-derived clots. Pathway enrichment analyses were then performed to understand biological mechanisms associated with each stroke subtype.

### Proteomic Analysis in the UK Biobank

This analysis was conducted within the UK Biobank, a population study of 503,317 adults aged 39 to 73 years at recruitment who were enrolled between 2006 and 2010 from across the United Kingdom. Referenced project application number is 58743. At baseline, participants provided blood samples, demographic and lifestyle information, and consented to long-term follow-up through linkage to national health records.(7) For the present study, we included participants with a documented ischemic stroke event and available plasma proteomic data. Participants were categorized according to the presence of ICD-10 codes for atrial fibrillation (I48) or carotid artery disease (I65). These groups were intended to represent ischemic stroke patients with coexisting AF or CA rather than adjudicated stroke etiologic subtypes. (8,9) Participants with missing protein measurements or concurrent diagnoses of both AF and carotid atherosclerosis were excluded. Proteomic profiling was performed using the Olink Explore 3072 platform, which measures 2,923 circulating proteins via proximity extension assays. Protein abundance values were normalized, rank-inverse transformed, and log2-scaled prior to analysis. Olink assay technology and analyses are described in detail elsewhere. (10,11) We fit linear regression models for each protein, with AF versus CA group status as the independent variable and protein concentration as the dependent variable. All models were adjusted for age, sex, ancestry principal components, body mass index, smoking status, hypertension, and diabetes. For each protein, we estimated the beta coefficient, standard error, and two-sided p-value. Multiple testing correction was applied using the Benjamini–Hochberg false discovery rate (FDR), and proteins with an FDR-adjusted p-value below 0.05 were considered statistically significant.

### Clot Single-cell Transcriptomics

To explore cellular expression patterns of candidate proteins, we analyzed single-cell RNA sequencing data from clots retrieved during mechanical thrombectomy for acute ischemic stroke at Yale New Haven Hospital. Clots were obtained from patients with ischemic stroke occurring in the setting of AF (n=7) or CA (n=7). Overall data processing, quality control, and annotation procedures have been described previously.(12) For the present study, we focused on genes encoding proteins that were significantly associated with stroke etiology in the UK Biobank proteomic analysis. Expression was assessed across annotated cell types using dot plots and average expression analyses, and qualitative comparisons were made across etiologic groups and cellular compartments to explore cell type–specific expression patterns and functional relevance of circulating protein markers. Non-parametric Kruskal–Wallis tests were used across annotated cell types, and Wilcoxon rank-sum testing was performed for focused comparisons when sufficient expression was detected.

### Mendelian Randomization

Proteins associated with AF versus CA group status in the proteomic analysis were further evaluated using two-sample Mendelian randomization (MR) analyses to assess associations between genetically predicted protein levels and ischemic stroke phenotypes. Cis-acting protein quantitative trait loci (cis-pQTLs) located within ±1 Mb of the encoding gene were used as instrumental variables. Variants were required to achieve genome-wide significance (P < 5×10⁻⁸) and were pruned for independence using an r² threshold of <0.001. MR analyses were conducted using publicly available summary statistics for any stroke (AS), any ischemic stroke (AIS), large artery stroke (LAS), cardioembolic stroke (CES) and small vessel stroke (SVS) from the GIGASTROKE consortium.(13) The primary MR method was inverse-variance weighted (IVW) regression. Sensitivity analyses included MR-Egger regression, weighted median estimation, and simple mode analyses to evaluate robustness to potential violations of instrumental variable assumptions. The number of cis-pQTL instruments ranged from 3 to 42 variants depending on the protein analyzed. Between-instrument heterogeneity was assessed using Cochran’s Q statistic, and horizontal pleiotropy was evaluated using the MR-Egger intercept test. Bidirectional MR analyses were additionally performed to assess potential reverse-direction associations between stroke phenotypes and protein levels.

### Pathway Enrichment

To explore biological pathways represented among proteins associated with AF or CA group status, we performed exploratory pathway enrichment analyses using the gseapy package.(14) Genes corresponding to proteins associated with AF– and CA–related stroke were analyzed separately. Enrichment was conducted against the 2022 Reactome and the 2021 Gene Ontology Biological Process databases, with statistical significance defined as an adjusted p-value (FDR) below 0.05.(15,16) Top enriched pathways were visualized as bar plots based on –log₁₀(FDR) to highlight biological processes and signaling pathways represented among identified protein sets.

### Software

Statistical analyses were performed using R version 4.2.1 and Python version 3.11.5 with the Seurat, multtest, and statsmodels packages. Mendelian randomization analyses were conducted using the Genal v1.3.2 framework.(17,18) Reporting followed the STROBE guidelines for observational studies and STROBE-MR recommendations for Mendelian randomization analyses.

## Results

### Proteomic Analysis in UK Biobank

We included 666 participants with ischemic stroke and available proteomic data, including 539 participants with atrial fibrillation (AF) and 127 with carotid atherosclerosis (CA). Mean age was slightly higher in the AF group compared with the CA group (63.3 vs 61.8 years, P=0.013). Sex distribution was similar across groups, with women representing 36.2% of participants in the AF group and 33.1% in the CA group (P=0.579) (**Table 1**).

**Table 1.** Demographic characteristics UKB cohort.

| **Variable** | **Overall**<br>**(n=666)** | **AF (n=539)** | **CA (n=127)** | **P-Value** |
| --- | --- | --- | --- | --- |
| **Age, mean (SD)** | 63.0 (5.8) | 63.3 (5.6) | 61.8 (6.3) | 0.013 |
| **Sex, n (%)** |  |  |  | 0.579 |
| - Female | 237 (35.6%) | 195 (36.2%) | 42 (33.1%) |  |
| **BMI, mean (SD)** | 29.2 (5.5) | 29.3 (5.5) | 28.7 (5.5) | 0.246 |
| **Hypertension, n (%)** |  |  |  | 0.895 |
| - Yes | 319 (47.9%) | 257 (47.7%) | 62 (48.8%) |  |
| **Diabetes, n (%)** |  |  |  | 0.955 |
| - Yes | 211 (31.7%) | 170 (31.5%) | 41 (32.3%) |  |
| **Smoking Status, n**<br>**(%)** |  |  |  | 0.033 |
| - Current | 113 (17.0%) | 82 (15.2%) | 31 (24.4%) |  |
| - Never | 252 (37.8%) | 214 (39.7%) | 38 (29.9%) |  |
| - Previous | 297 (44.6%) | 239 (44.3%) | 58 (45.7%) |  |
| - Prefer not to answer | 4 (0.6%) | 4 (0.7%) | 0 (0.0%) |  |

In unadjusted analyses, NT-proBNP, NPPB, and ACP5 were significantly higher among ischemic stroke participants with AF, whereas F7, F10, ANGPT2, APCS, PROS1, and LARP1 were higher among participants with CA (all FDR < 0.05). The strongest associations were observed for NT-proBNP (β = –1.23, P = 1.9×10⁻¹¹) and NPPB (β = – 1.18, P = 2.6×10⁻⁹) (**Figure 2**, Supplementary Table 1). In adjusted models, twelve circulating proteins differed significantly between AF and CA groups after correction for multiple testing (all FDR <0.05). Compared with the CA group, the AF group demonstrated higher plasma levels of NTproBNP (β = –1.15, SE 0.18, FDR = 1×10⁻⁶), NPPB (β = –1.13, SE 0.20, FDR = 1.9×10⁻⁵), and ACP5 (β = –0.21, SE 0.04, FDR = 8.7×10⁻⁵). Conversely, lower levels of APCS (β = 0.20, SE 0.05, FDR = 1.2×10⁻²), ANGPT2 (β = 0.20, SE 0.05, FDR = 2.6×10⁻²), PAMR1 (β = 0.13, SE 0.03, FDR = 1.2×10⁻²), PRCP (β = 0.15, SE 0.04, FDR = 1.4×10⁻²), PROS1 (β = 0.17, SE 0.04, FDR = 1.4×10⁻²), LARP1 (β = 0.25, SE 0.06, FDR = 1.9×10⁻²), F7 (β = 0.25, SE 0.05, FDR = 4.5×10⁻⁴), F10 (β = 0.20, SE 0.05, FDR = 1.2×10⁻²), and LEO1 (β = 0.11, SE 0.03, FDR = 4.7×10⁻²). were observed in the AF group relative to the CA group. (**Figure 2**, **Table 2**) NTproBNP and NPPB remained significant after Bonferroni correction.

**Figure 2.**
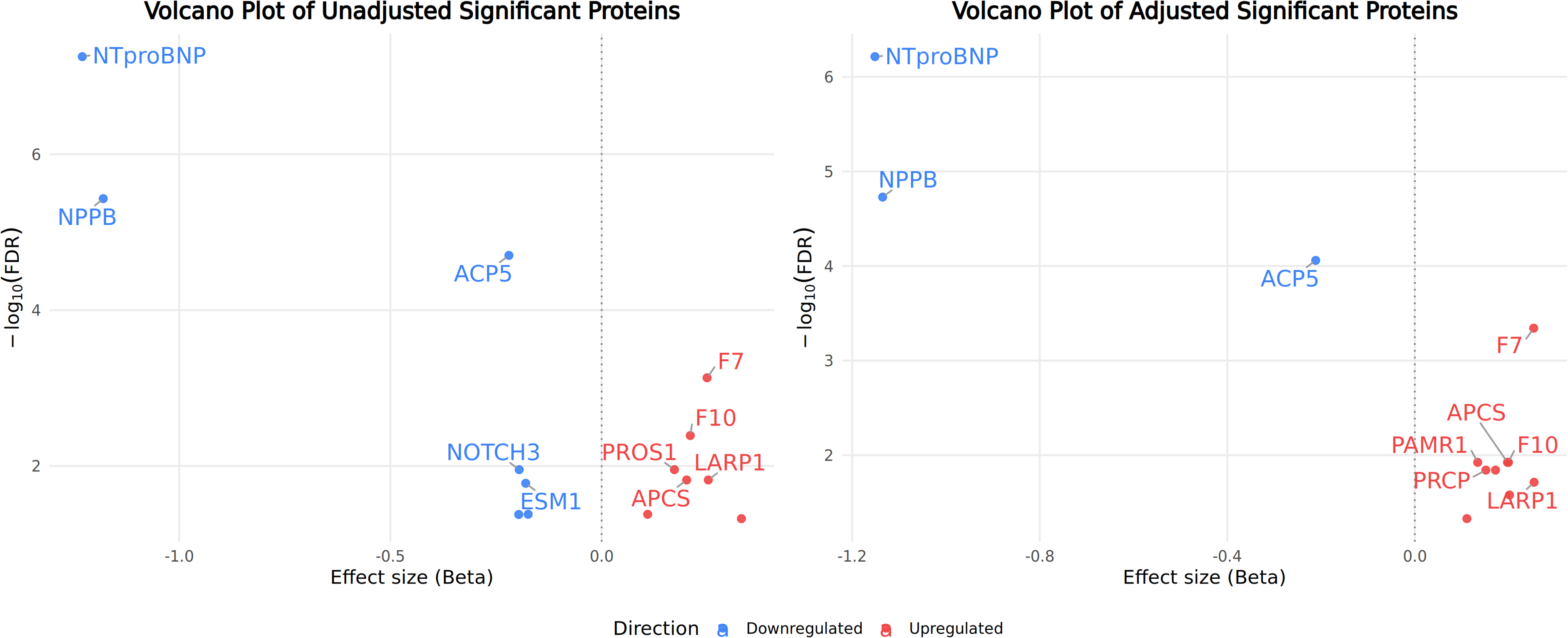
Volcano plots of proteomic associations between cardioembolic and carotid atherosclerotic stroke (unadjusted and adjusted models).

**Table 2.** Differentially Expressed Plasma Proteins Between Cardioembolic and Carotid Atherosclerotic Stroke in the UK Biobank (Adjusted Models).

| Beta | SE | P | FDR | Bonferroni |
| --- | --- | --- | --- | --- |
| NTproBNP | -1.15141 | 0.178233 | 2.09E-10 | 6.11E-07 |
| NPPB | -1.13495 | 0.196795 | 1.28E-08 | 1.87E-05 |
| ACP5 | -0.21141 | 0.039077 | 8.97E-08 | 8.73E-05 |
| F7 | 0.253564 | 0.05035 | 6.22E-07 | 4.54E-04 |
| APCS | 0.197641 | 0.046495 | 2.53E-05 | 1.19E-02 |
| PAMR1 | 0.134136 | 0.031435 | 2.29E-05 | 1.19E-02 |
| F10 | 0.199647 | 0.047289 | 2.86E-05 | 1.19E-02 |
| PRCP | 0.151502 | 0.036843 | 4.44E-05 | 1.44E-02 |
| PROS1 | 0.172059 | 0.041714 | 4.32E-05 | 1.44E-02 |
| LARP1 | 0.254406 | 0.063254 | 6.64E-05 | 1.94E-02 |
| ANGPT2 | 0.202099 | 0.051597 | 9.95E-05 | 2.64E-02 |
| LEO1 | 0.111147 | 0.029598 | 1.93E-04 | 4.69E-02 |

### Clot Single-cell Transcriptomics

Exploratory single-cell RNA sequencing analyses of thrombectomy clots demonstrated differential expression for several candidate proteins identified in the proteomic analysis. ACP5 expression higher in clots from patients with AF compared with CA (mean normalized expression 0.80 vs 0.60, *P*<0.001), while PRCP (0.32 vs 0.28, *P*<0.001) and LARP1 (0.58 vs 0.47, *P*<0.001) demonstrated relatively higher expression in clots from patients with CA. PROS1 expression did not significantly differ between groups (0.03 vs 0.026, *P*=0.07) (**Figure 3, Supplementary Table 2).** When examined across annotated cell types, ACP5 expression was greatest in macrophages, with average expression exceeding 4.0, substantially higher than in monocytes (0.76), dendritic cells (0.32), or other immune populations. PRCP and LARP1 demonstrated broader expression across macrophages, dendritic cells, and T cell subsets. Global non-parametric testing confirmed significant differences across cell populations for ACP5, PRCP, and LARP1 (Kruskal–Wallis *P*<0.001 for all). Focused comparisons with Wilcoxon rank-sum testing further supported macrophages as the principal source of ACP5 in AF-related clots. (**Figure 4, Supplementary Table 3)**

**Figure 3.**
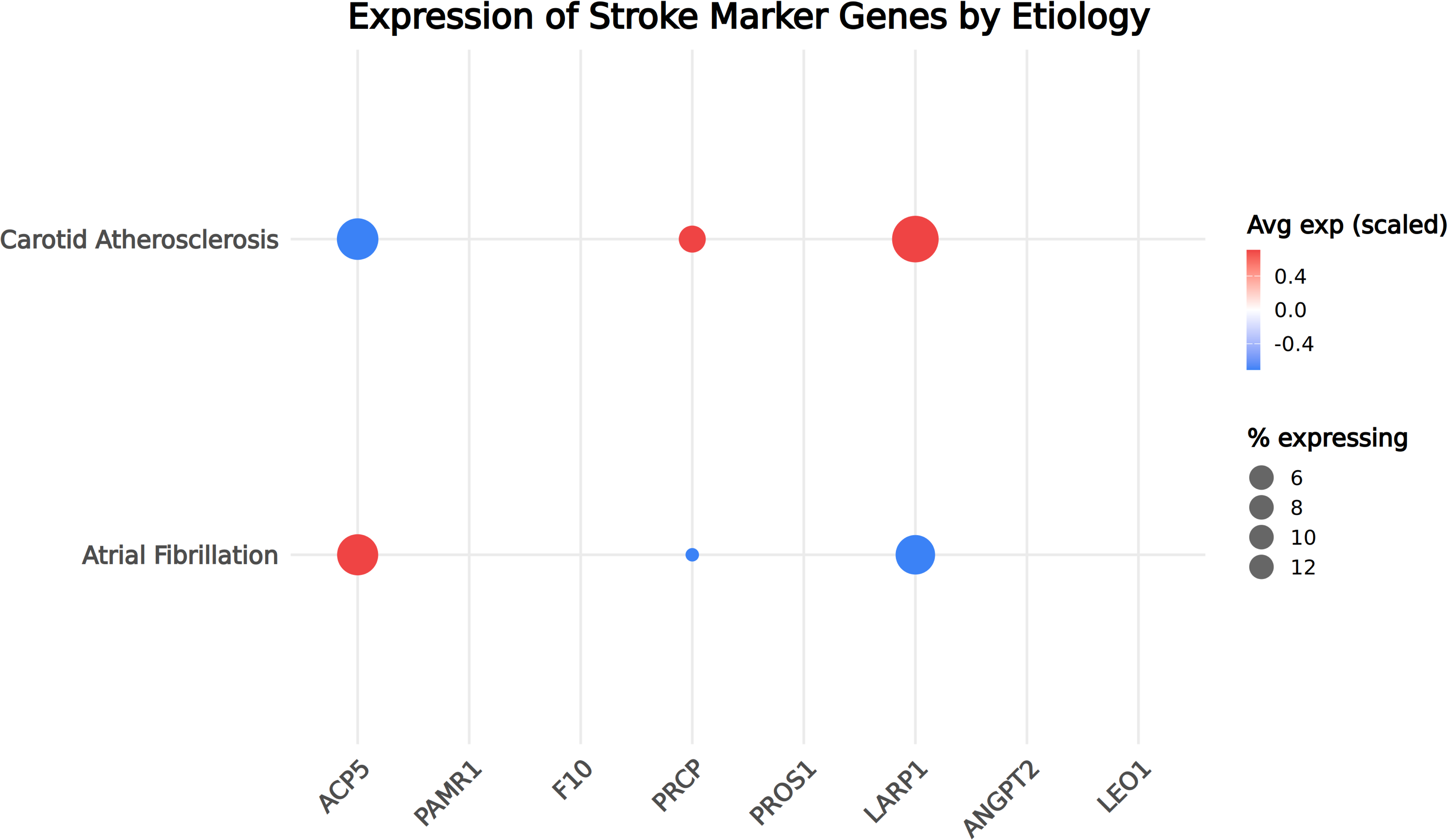
Single-cell RNA sequencing of retrieved thrombi showing differential expression of candidate proteins across stroke etiologies (dot plots).

**Figure 4.**
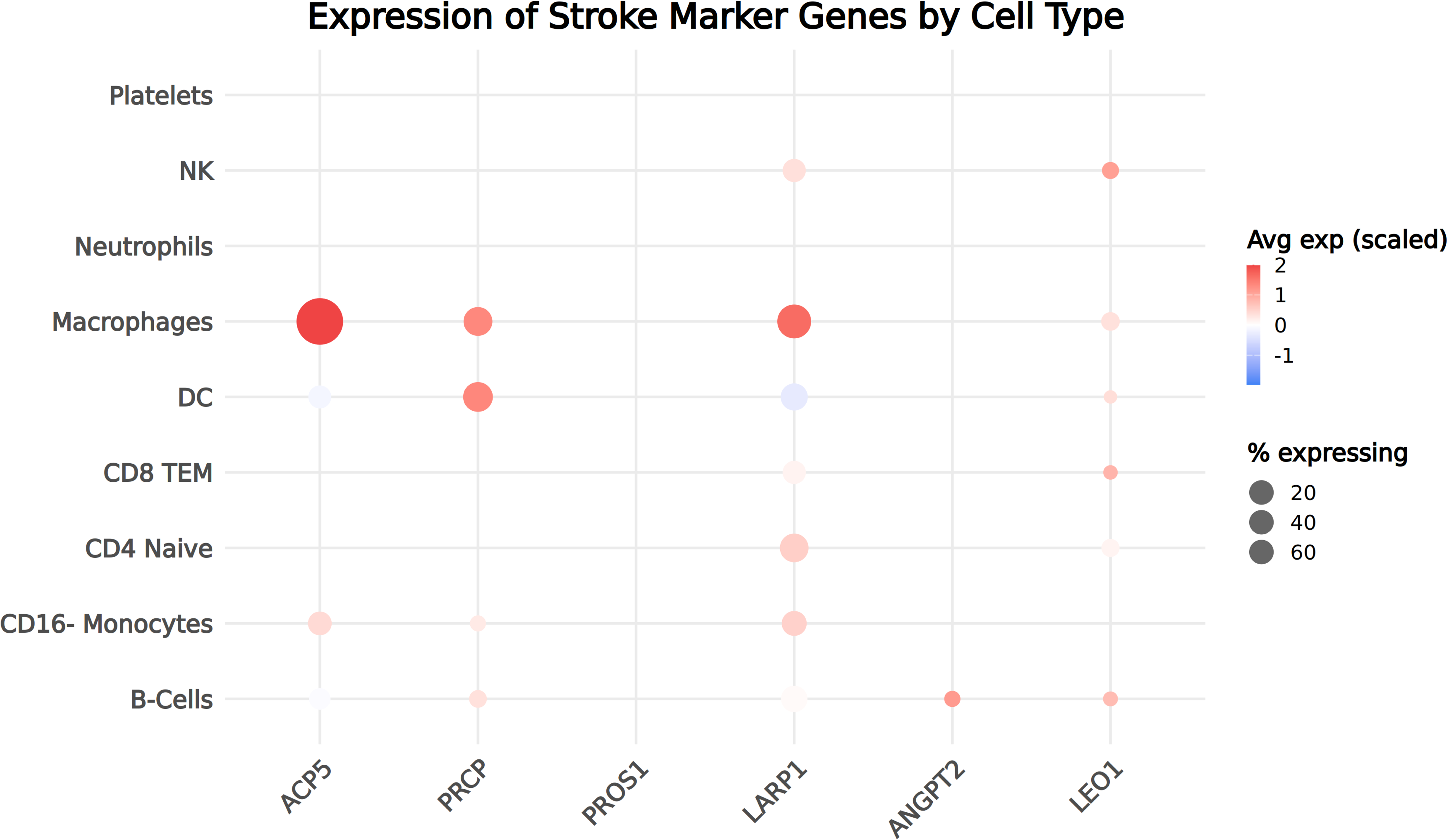
Single-cell RNA sequencing of retrieved thrombi showing differential expression of candidate proteins across clots cell types (dot plots).

### Mendelian Randomization Analyses

Fixed-effects inverse variance–weighted MR analyses demonstrated associations between genetically predicted protein levels and ischemic stroke phenotypes. Genetically predicted NT-proBNP levels were associated with cardioembolic stroke (CES) (OR 1.26, 95% CI 1.11–1.42, p = 2.1×10⁻⁴). APCS (OR 1.08, 95% CI 1.01–1.16, p = 0.022), ANGPT2 (OR 1.07, 95% CI 1.01–1.13, p = 0.020) were significantly associated with any ischemic stroke (AIS), and (OR 1.06, 95% CI 1.01–1.10, p = 0.017) were associated with any ischemic stroke (AIS). ANGPT2 was additionally associated with small vessel stroke (SVS) (OR 1.17, 95% CI 1.01–1.35, p = 0.036). NPPB (OR 1.48, 95% CI 1.15–1.90, p = 0.002) and PAMR1 (OR 1.11, 95% CI 1.02–1.20, p = 0.018) were associated with large artery stroke (LAS). Results were directionally consistent across sensitivity analyses, including weighted median and MR-Egger methods. No significant evidence of horizontal pleiotropy was identified. In bidirectional analyses, there was no evidence that ischemic stroke phenotypes genetically influenced circulating protein levels. **(Supplementary Table 4)**

### Pathway Enrichment

Exploratory pathway enrichment analyses suggested enrichment of natriuretic peptide signaling and riboflavin metabolism pathways among proteins associated with the AF group. Proteins associated with the CA group were enriched for coagulation-related and gamma-carboxylation pathways, including fibrin clot formation and gamma-carboxylation of protein precursors (**Figure 5, Supplementary Table 5-6).**

**Figure 5.**
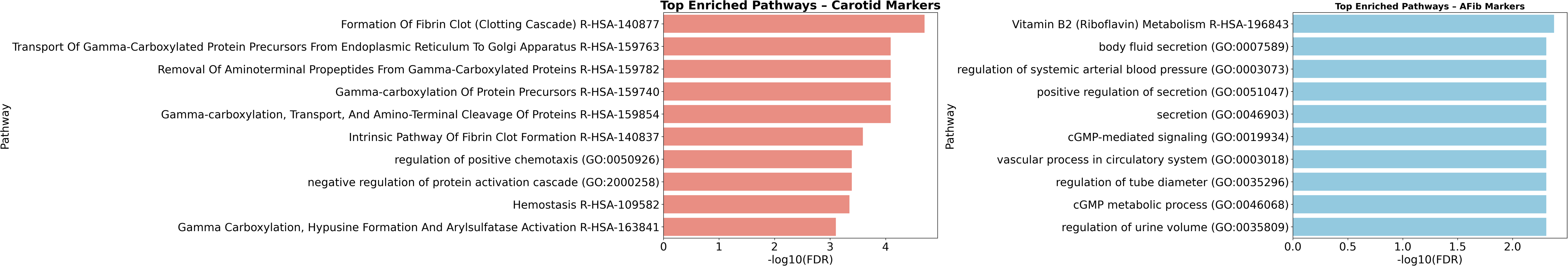
Pathway enrichment analyses highlighting distinct biological processes in cardioembolic versus carotid atherosclerotic stroke (bar plots).

## Discussion

In this multimodal analysis using complementary proteomic, transcriptomic, and genetic datasets, we identified differing molecular profiles among ischemic stroke patients with atrial fibrillation (AF) versus carotid atherosclerosis (CA). Several observed associations were consistent across orthogonal analytical approaches, including circulating plasma proteomics, exploratory clot single-cell transcriptomic analyses, and Mendelian randomization studies. Together, these findings support the presence of differentially expressed pathways in ischemic stroke occurring in the setting of AF or CA.

Confirming the validity of the results obtained with this analytical approach, several of the identified proteins correspond to biomarkers previously linked to AF and vascular disease. NT-proBNP demonstrated the strongest association with the AF group, consistent with extensive prior literature linking natriuretic peptides with atrial remodeling, arrhythmia burden, and cardioembolic stroke risk.(19,20) Similarly, higher levels of coagulation-related proteins, including factor VII among participants with CA, are biologically consistent with activation of thrombotic pathways in atherosclerotic vascular disease.(21–23)

Beyond established biomarkers, our analysis identified several novel proteins that warrant further investigation as candidate biomarkers. Among participants with AF, ACP5 emerged as a notable signal across both proteomic and exploratory transcriptomic analyses. ACP5 is a macrophage-associated enzyme previously implicated in inflammatory and fibrotic pathways.(24–26) Although the present study cannot definitively establish mechanistic causality or cellular origin, the observed association raises the possibility that inflammatory and profibrotic processes may contribute to the molecular profile observed among ischemic stroke patients with AF.

Among participants with CA, several proteins associated with endothelial dysfunction, coagulation, and vascular inflammation demonstrated differential expression patterns. ANGPT2, a regulator of endothelial activation and vascular remodeling, has previously been linked to plaque instability and atherosclerotic disease progression.(27,28) PRCP, which participates in the renin–angiotensin and kallikrein–kinin systems, has been associated with endothelial dysfunction and thrombosis in experimental models.(29–31) Additional proteins including APCS, PROS1, and F10 also demonstrated associations with the CA group, although their biological roles in ischemic stroke remain incompletely understood and likely context dependent.(32–36)

We also identified several less well-characterized intracellular regulators, including LARP1, LEO1, and PAMR1, that have not been extensively studied in cerebrovascular disease. These proteins are involved in cellular signaling, transcriptional regulation, and tissue remodeling pathways.(36–38) Although exploratory, these findings may provide direction for future mechanistic studies evaluating vascular and thromboinflammatory biology in ischemic stroke.

The clinical implications of these findings should be interpreted cautiously. Although these molecular profiles are not sufficient for etiologic classification, they may contribute to improved biological characterization of ischemic stroke populations and help identify candidate pathways for future translational investigation. In particular, proteins such as ACP5 and ANGPT2 may represent potential targets for future experimental studies focused on atrial remodeling, vascular inflammation, or plaque instability.

Several limitations merit consideration. First, classification of participants was based on ICD coding for AF and carotid artery disease rather than adjudicated stroke mechanisms, and these groups should therefore be interpreted as ischemic stroke patients with coexisting AF or CA rather than definitive cardioembolic or atheroembolic stroke subtypes. Second, the UK Biobank proteomic measurements were obtained at baseline, often years before stroke occurrence, and therefore likely reflect chronic biological risk states rather than acute stroke biology. Third, the clot single-cell transcriptomic analyses were derived from a small thrombectomy cohort and should be considered exploratory. Cell-level differential expression analyses may also be influenced by pseudoreplication and clinical heterogeneity. Fourth, the independent datasets used in this study represent distinct biological contexts and were not integrated at the individual patient level. Finally, although Mendelian randomization analyses supported several protein–stroke associations, these findings should not be interpreted as definitive evidence of causality.

## Conclusion

In conclusion, complementary proteomic, transcriptomic, and genetic analyses identified differing molecular patterns among ischemic stroke patients with AF versus CA. These findings support previously recognized biomarkers while also identifying additional candidate pathways that may contribute to differing vascular and thromboinflammatory profiles. Further validation in larger, clinically adjudicated, and longitudinal cohorts will be necessary to clarify the biological and translational relevance of these observations.

